# On the relationship between neuronal heterogeneity and entropy

**DOI:** 10.1101/2025.06.07.658464

**Authors:** Taufik A Valiante

## Abstract

Neuronal size has often been used to explain the “superiority” of the human brain. By deriving the Shannon entropy of different statistical distributions, we show that the entropy of different distributions is solely accounted for by the variance of the distribution. We use total dendritic length of neurons from different species as an example to show that entropy computed from TDL distributions has nothing to do with the absolute size of neurons, but the increased variance seen in the larger neurons.

## Introduction

Computational models have demonstrated that intrinsic biophysical heterogeneity within a population of neurons leads to greater coding efficiency and higher information content than homogeneous populations ***Shamir and Sompolinsky (2006)***; ***Mejias and Longtin (2012)***; ***Chelaru and Dragoi (2008)***; ***Padmanabhan and Urban (2010)***; ***Tripathy et al. (2013)***.

While such approaches provide specific examples of the relationship between diversity and information coding, we here sought to provide derivations of Shannon Entropy which are independent of network architecture, and statistical properties of experimentally recorded neurons.

Our results provide an immediate understanding of why larger neurons of the human brain confer an information benefit independent of numbers and size alone. Our results also provide a conceptual framework for understanding how declines in heterogeneity may result in information poor states of the brain resulting in cognitive and behavioral declines.

## Results

We briefly enumerate the results from the Appendix where we derive the Shannon entropy of a number of exemplary distributions.

### Uniform Distribution

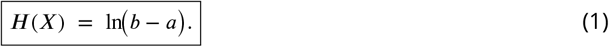

Where *H*(*X*) is the Shannon entropy ***Shannon (1948)*** of a randomly distributed variable *X* sampled from a uniform distribution and a and b represent upper and lower bounds over which *X* is sampled.

### Normal Distribution

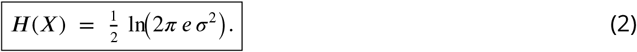

Where *σ*2 represents the variance of the normally distributed observations.

### Log-Normal Distribution

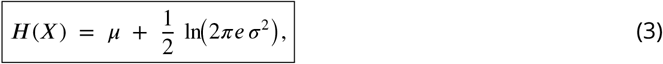

Where *μ* and *σ*^2^ are mean and the variance of the underlying normal distribution.

From equations 1 and 2 it can immediately seen that entropy is solely dependent on the spread of the data, and not the mean. For the normal distribution *H*(*X*) is *solely* dependent on *σ*^2^ while for the uniform distribution it is the range of the data. These results confirm the intuition underlying the concept of entropy, whereby is is solely dependent on the variability/uncertainty of *X* and nothing to do with where *X* is centered.

Interestingly however this is not the case for the log-normal distribution, a distribution that characterizes a large number of natural occurring phenomena and cell and circuit features of the brain ***Buzsáki and Mizuseki (2014)***. From 3 it can be seen that *H*(*X*) depends linearly on the mean and logarithmically on the variance of the underlying normal distribution.

In **Figure 1A** we plot the relationship between *H*[*X*] and the *E*[*X*] using Equation 3. For this computation we use a variance (*σ*^2^) computed from Equation 4 of the underlying normal distribution for mouse total dendritic length (TDL) data from ***Mertens et al. (2024)*** with a mean and SD of 3.9 and 0.6 respectively. We then plot the entropy for different species reported in ***Mertens et al. (2024)*** and from ***Altemus et al. (2005)*** for monkey on the same axis assuming that the TDL observation were drawn from either a normal distribution or log-normal distribution. Plotting the data in this way, we can immediately see that information scales as the size of the neurons increases. From this we might conclude that neuronal size drives the increase in information content. However interestingly if we compute *H*[*X*] assuming the samples are drawn from a normal distribution (where we know *H*[*X*] is independent of the mean), we can see that there is little difference in *H*[*X*] under these two different cases - *μ* contributes almost nothing to the information content for the log-normal distribution. However, this conclusion seems odd, since there is clearly a strong correlation between *H*[*X*] and *E*[*X*]. How can we reconcile this? From the derivation in the appendix and with a little rearrangement we obtain Equation 4 where we can see *Var*[*X*] scales quadratically with *E*[*X*].

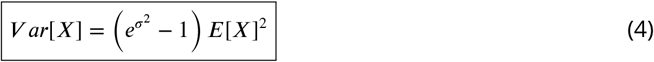

Where *E*[*X*] and *Var*[*X*] are the observed mean and variance of experimental observations *X*, and *σ*^2^ is the variance of the underlying normal distribution.

**Figure 1.**
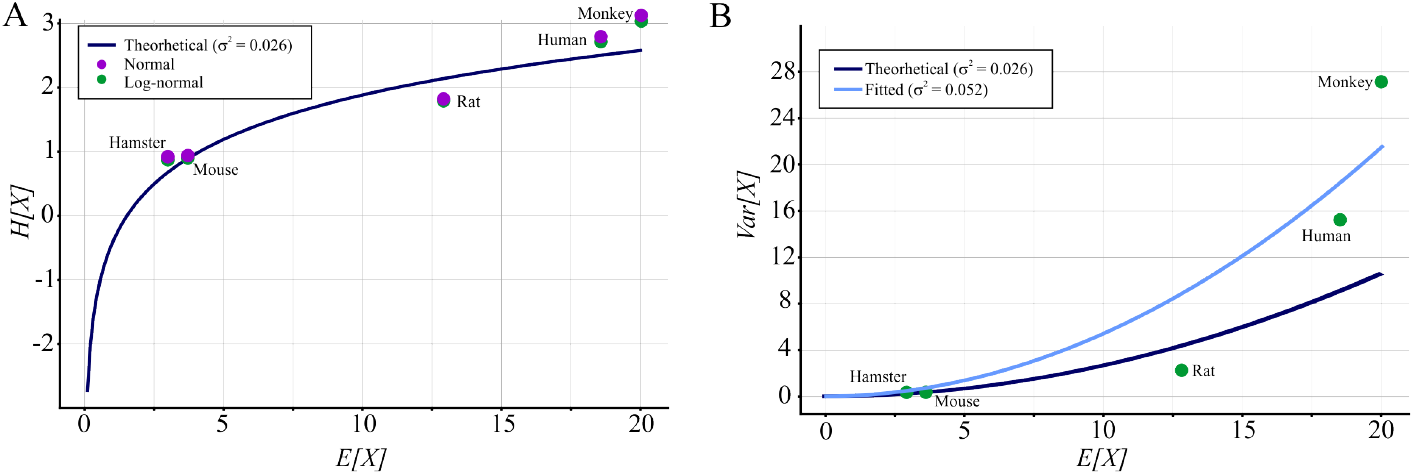
The inter-relationship of the expected value of a log-normally distributed variable and its variance. **A)** *H*[*X*] is plotted using 3 with *σ* representing the standard deviation of the underlying normal distribution. *E*[*X*] is the expected value of the log-normally distributed variable *X*, the range of which was chosen to cover measurements of total dendritic length (TDL) for different species reported in ***Mertens et al. (2024)***. *σ* = 0.026 was obtained from equation 4 using mouse values (*E*[*X*] = 3.7; *Var*[*X*] = 0.6) ***Mertens et al. (2024)***. Monkey data was obtained from ***Altemus et al. (2005)***. **B)** Relationship of the measured variance (*Var*[*X*]) of a log-normally distributed random variable and its mean (*E*[*X*]) plotted using equation 4 for two values of *σ*^2^. The first is derived from mouse data (as in A above), and the second from a non-linear curve fit of equation 4 to the experimental data

This is because the location of the mean of the underlying normally distribution changes the shape of the distribution when log transformed. Thus the variance of the log-normally distributed variance is dependent on the mean. The mean thus drives variance. This can be visualized in **Figure 1B** where *V ar*[*X*] (experimentally measured variance) is plotted as a function of their *E*[*X*] (measured experimental means). As in **Figure1A** we use the *σ*^2^ for mice TDL. Plotted in this way if the primate was on this theoretical curve we could conclude that primate neurons are just scaled versions of mouse neurons, which of course we know is not the case ***Benavides-Piccione et al. (2020)***. Also there is an additional ‘boost’ of variability for human and monkey neurons that can not be attributed to a simple scaling up of mouse neurons. This additional variability then contributes to the boost in *H*[*X*] seen in **Figure 1A** for human and monkey neurons.

Alternatively we can find the *σ*^2^ that best “fits” the data (Figure 1B) - in other words to see if there is a *σ*^2^ of an underlying normal distribution that well describes our dataset. Interestingly despite providing a reasonable fit for hamster and mouse neurons the larger *σ*^2^ of 0.052 poorly fits (as did *σ*^2^ = 0.026) the human and monkey data. It is of course surprising that monkey neurons are larger and have the largest *V ar*[*X*] (more diverse). Our recent work provides a plausible explanation for this seeming discrepancy. We suggest that the human variance is underestimated, since the sample are taken from epilepsy patients***Mertens et al. (2024)***, a condition we have shown to associated with decreased variance in biophysical properties ***Rich et al. (2022)***; ***Chameh et al. (2023)***. Our work has shown that the intrinsic biophysical properties of human L5 cortical neurons are less diverse when sampled from the epileptogenic zone (area of the brain that generates seizures) of epilepsy patients ***Rich et al. (2022)***; ***Chameh et al. (2023)*** when compared to regions of the brain not involved in generating seizures. We have also shown that human cortical neurons, and dentate granule neurons of the hippocampus from humans with epilepsy, show a decrease in transcriptomic diversity ***Chameh et al. (2023)*** - paralleling our electrophysiological findings ***Rich et al. (2022)***. Thus it is very likely that the human data point is an underestimation fo the true *Var*[*X*] in TDL of hippocampal CA1 neurons. Interestingly such decreases in morphological diversity can be observed in slice culture (see Figure of ***Lee et al. (2023)***) - another “pathological” state of the brain.

## Discussion

Here we demonstrate a fundamental relationship between the variance of a probability distribution and its entropy. We show that the Shannon Entropy of a randomly distributed variable is always related to its variance, the specific form of this relationship however differing for different types of distributions. The log-normal distribution is unique when compared to the uniform and normal distributions, where it variance depends on the mean as well. This is because the mean of the underlying normal distribution alters the shape of the log-transformed distribution which then alters the variance. Specifically the observed variance scales quadratically with the expected mean of the data, and thus in the context of neuronal TDLs, larger neurons will have scaled up variances due to the larger size. However although some of the variability might be accounted for by this scaling, there seems to be additional unaccounted variance that “boost” the morphological diversity of monkey and human neurons.

Although we have chosen to show this scaling for a specific feature of neurons - TDL - it is likely to hold for a number of other morphological and biophysical properties as a large number of features of the brain can be well described by the log-normal distribution***Buzsáki and Mizuseki (2014)***.

Overall our findings here provide evidence for a general relationship between a distributions variance and its coding capacity. Specifically for neural systems, information coding capacity is better thought of as a feature that scales with the variance of constituent elements rather than their magnitude.

## Methods and Materials

Curve fitting was performed in Pyhton using the curve-fit function found in the optimize package of scipy. It uses a non-linear least squares approach employing the the Levenberg-Marquardt algorithm.

## Appendix 1

### Normal distribution

Here we derive the (differential) Shannon entropy of a continuous random variable *X* which follows a normal (Gaussian) distribution with mean *μ* and variance *σ*^2^. Recall that the probability density function (pdf) of a normal random variable *X ∼* 𝒩 (*μ, σ*^2^) is given by

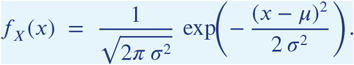

We aim to compute its differential entropy,

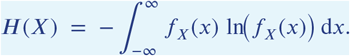

Step 1: Express the Logarithm of the PDF

First, we write the logarithm of the density *f*_*X*_(*x*):

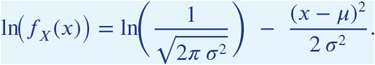

Since 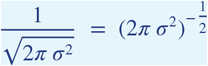, it follows that

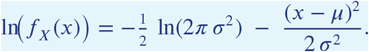

Step 2: Plug into the Definition of Entropy

The differential entropy is

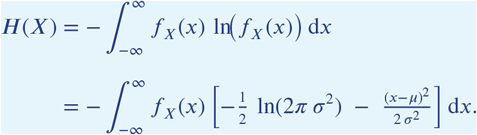

Distribute the negative sign:

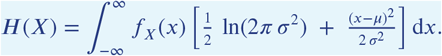

Step 3: Separate the Integral into Two Parts

We can split the integral into two parts:

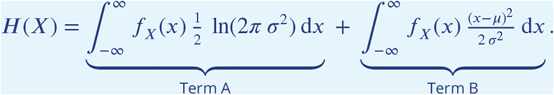

Term A

Since 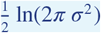 ln(2*π σ*^2^) does not depend on *x*, it comes out of the integral:

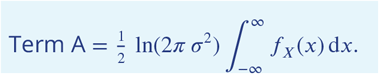

Because *f*_*X*_(*x*) is a pdf, its integral over the entire real line is 1:

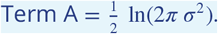

Term B

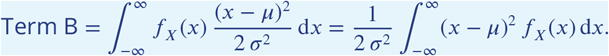

But the integral 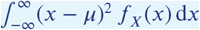 is precisely the variance *σ*^2^. Hence,

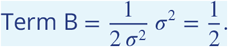

Step 4: Combine Both Terms

Putting the two terms together,

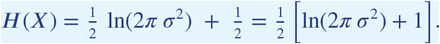

Often, we rewrite this by factoring out 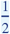 and using the property ln(*a*) + 1 = ln(*e a*), giving

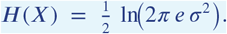

#### Final Result

Hence, the differential entropy of a normal random variable *X ∼* 𝒩 (*μ, σ*^2^) is

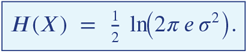

### Uniform distribution

A random variable *X* is said to be **uniformly distributed** on the interval [*a, b*] (with *a* < *b*) if its probability density function (pdf) is

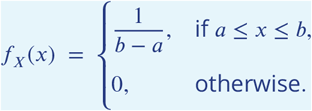

We seek to compute its **entropy**:

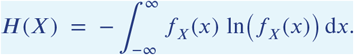

Step 1: Plug in the Definition of Entropy

Since *f*_*X*_(*x*) is nonzero only for *x* ∈ [*a, b*], the integral reduces to

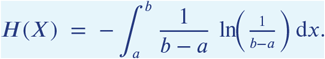

Note that 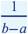 is a constant with respect to x.

Step 2: Bring Out Constants

We can factor out – ln 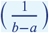 and 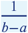 from the integral:

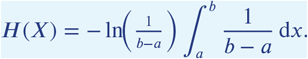

Step 3: Evaluate the Remaining Integral

The integral of the constant 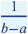 from to is simply 1:

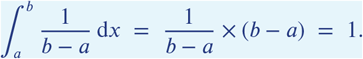

Hence,

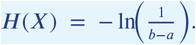

Step 4: Simplify the Logarithm

We know that 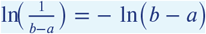. Thus,

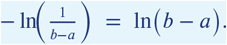

Therefore,

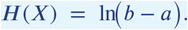

#### Final Result

Hence, the entropy of a uniform random variable *X ∼* Uniform(*a, b*) is:

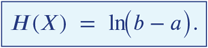

### Log-normal distribution

A random variable *X* is said to have a **lognormal distribution** if ln(*X*) (the natural logarithm of *X*) follows a normal distribution with parameters *μ* and *σ*^2^. Equivalently, if *Z ∼* 𝒩 (*μ, σ*^2^), then

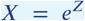

is lognormally distributed. The probability density function (pdf) of *X* is

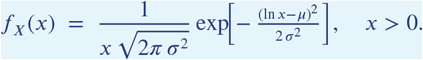

We seek to compute the **differential entropy**

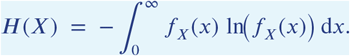

Step 1: Express ln(*f*_*X*_(*x*))

Write out the logarithm of *f*_*X*_(*x*):

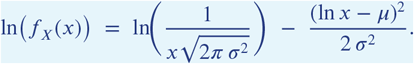

Separating terms,

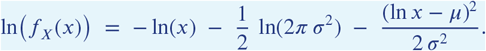

Define

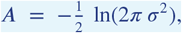

so that

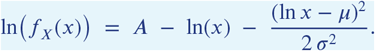

Step 2: Plug into the Definition of Entropy

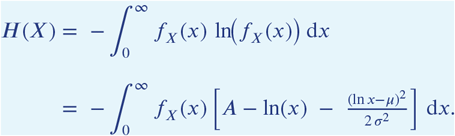

Distribute the minus sign inside:

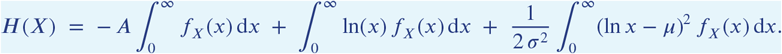

Step 3: Evaluate Each Integral Separately

(a) 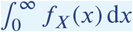

Because *f*_*X*_(*x*) is a pdf on (0, ∞),

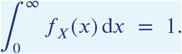

Hence

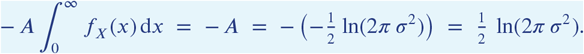

(b) 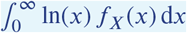

By definition of the lognormal distribution, if *X* = *e*^*Z*^ with *Z ∼* 𝒩 (*μ, σ*^2^), then

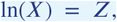

which has mean *μ*. Therefore

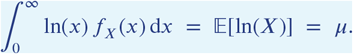

(c) 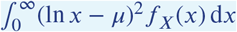

Similarly, ln(*X*) has variance *σ*^2^. Hence

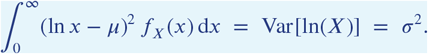

Thus

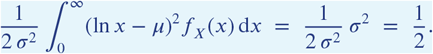

Step 4: Combine All Terms

Summing these contributions, we get

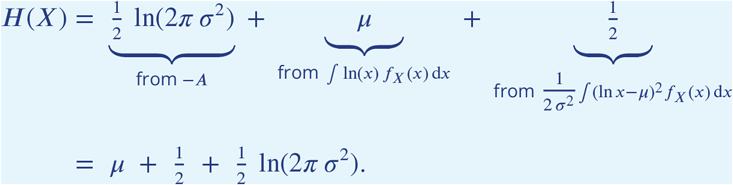

We can rewrite 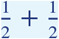 as 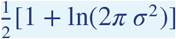. Noting that 1 = ln(*e*), we get

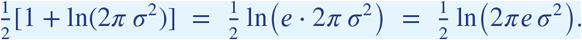

Hence the final expression is

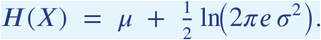

#### Final Result

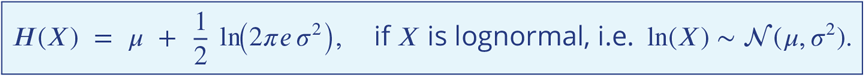

### Variance of underling normal distribution of a Log-normally distributed variable

To derive *σ*^2^, the variance of the underlying normal distribution, we start with the variance formula of a log-normal distribution:

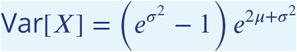

Divide through by 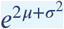:

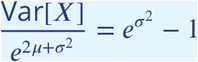

Rearrange to isolate 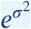:

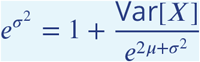

Substitute 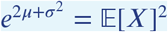 (from the relationship between mean and variance):

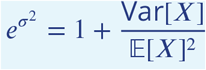

Take the natural logarithm of both sides:

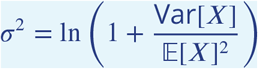

Thus, the variance of the underlying normal distribution is:

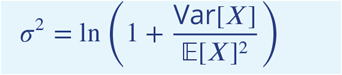

